# Anchors for Time, Distance, and Magnitude in Virtual Movements

**DOI:** 10.1101/2022.09.12.507649

**Authors:** Keri Anne Gladhill, Eva Marie Robinson, Candice Stanfield-Wiswall, Farah Bader, Martin Wiener

## Abstract

In order to navigate through the environment, humans must be able to measure both the distance traveled in space, and the interval covered in time. Yet, how these two dimensions are computed and interact across neural systems remains unknown. One possibility is that subjects measure how far and how long they have traveled relative to a known reference point, or anchor. To measure this, we had human participants (n=24) perform a distance estimation task in a virtual environment in which they were cued to attend to either the spatial or temporal interval traveled, while responses were measured with multiband fMRI. We observed that both dimensions evoked similar frontoparietal networks, yet with a striking rostrocaudal dissociation between temporal and spatial estimation. Multivariate classifiers trained on each dimension were further able to predict the temporal or spatial interval traveled, with centers of activation within the supplementary motor area (SMA) and retrosplenial cortex (RSC) for time and space, respectively. Further, a cross-classification approach revealed the right supramarginal gyrus (SMG) and occipital place area (OPA) as regions capable of decoding the general magnitude of the traveled distance. Altogether, our findings suggest the brain uses separate systems for tracking spatial and temporal distances, which are combined together along with amodal estimates.

## Introduction

The perception and production of both temporal and spatial features of the environment is necessary for mobile organisms to interact with and navigate through the world. However, it is not yet fully understood whether these dimensions rely on shared or differential neural circuits as the current literature supports both possibilities (1; 2; 3; 4; 5). These divergent findings could be due to the natural correlation of both temporal and spatial magnitudes in that longer distances take more time and shorter distances take less time, making it difficult for humans to attend to spatial information without also taking into account temporal information (6). In fact, brain areas such as the prefrontal cortex and right parietal cortex have been implicated in different types of magnitude processing; specifically those of time and space (4). Additional brain areas that have been implicated in the estimation and encoding of temporal duration during functional magnetic resonance imaging (fMRI) include the right dorsolateral prefrontal cortex, supplementary motor area (SMA), basal ganglia, and inferior frontal gyrus (7; 8). Other studies suggest a neural disassociation between temporal and spatial processing; specifically, spatial and distance tasks activate more posterior regions such as the parahippocampus, anterior hippocampus, and retrosplenial complex (9; 10; 11; 12), whereas temporal tasks exclusively activate the SMA (13). A meta-analysis of neuroimaging studies of this type revealed partially overlapping regions of activation likelihood across these various brain regions; specifically, the SMA, right inferior parietal lobe, and right inferior frontal gyrus (14).

More recently studies have demonstrated the ability to disentangle time and space from one another by using virtual reality (VR) environments (15; 16; 17; 18; 19; 3). Specifically, participants performed a virtual reproduction task in which they moved forward virtually in the virtual environment (VE) for an unknown distance or length of time (estimation) then reproduced the distance or time by moving forward in the VE (reproduction). Importantly, in order to dissociate time from space, the speed at which participants moved varied between the estimation and reproduction phases (15; 17; 18; 20). Therefore, participants were either to pay attention to the amount of time they moved and reproduce that time or pay attention to the amount of distance they moved and reproduce that distance. This has previously been tested behaviorally (15; 18) as well as using electroencephalogram (EEG) (21) in all cases both space and time exhibited central tendency effects in which short intervals or distances wereoverestimated and long intervals or distance were underestimated (22).

Importantly, the EEG study (21) focused on the event-related potential (ERP) known as the CNV (contingent negative variation) which has been suggested to be driven by the SMA and corticothalamic circuitry (23). The CNV has additionally been shown to have a more negative amplitude for longer duration estimates in a variety of timing tasks (24; 25; 26; 27). However, other studies suggest that the CNV may instead reflect a more general neural signature for the processing of different magnitudes (28; 29). Furthermore, the CNV has demonstrated a general sense of expectation in which the probability of an upcoming stimuli occurring in time is encoded, which would be critical during spatial navigation (30; 31; 32; 33). The findings from this EEG study revealed a dissociation within the initial negative deflection of the CNV, or iCNV, between tasks during reproduction in that, the amplitude for temporal estimates covaried with the length of the interval participants planned to reproduce; however, there was no difference in the iCNV for spatial estimates. This finding suggests that there exist differential processing modes for the reproduction of temporal intervals compared to spatial distances.

In the current study, we sought to further disassociate the neural mechanisms of time and space perception by having participants perform the same exact task as used in Robinson et al (2021) while undergoing fMRI.

## Results

To interrogate time and distance, we employed a virtual walking task as used previously by our lab and others (17; 15; 21; 18). In this task, subjects are placed in a virtual environment (VE) consisting of an open world landscape with distant rocks and a target sphere on the horizon (Figure 1). Subjects are required to walk forward while estimating either the time or distance of their movement (estimation phase), and then must reproduce that same magnitude again (reproduction phase). Crucially, the speed with which subjects move was randomly changed between estimation and reproduction phases to ensure that subjects could not use the secondary dimension (i.e. time) when estimating the primary dimension (i.e. distance). All subjects received feedback for reproducing stimuli within an adaptive window centered on a proportion of the base interval. Similar to previous reports, we observed that subjects exhibited a central tendency effect, in which reproduced estimates gravitated to the mean of the stimulus set. Notably, this effect was larger for time than for distance [t(15)=-3.078, *p*=0.008, Cohen’s *d* = 0.769], and was also correlated between dimensions [*r* = 0.7, *p*=0.003], suggesting a common mechanism for reproducing magnitudes (5). In contrast, when examining the variability of reproduced estimates, we observed no difference between time and distance [*F* (1,15)=0.467, *p*=0.505], nor any interaction with the tested interval [*F* (6,90)=1.156, *p*=0.337], and no correlation between the two [*r =* -0.048, *p* = 0.86] (Figure 1).

**Figure 1:**
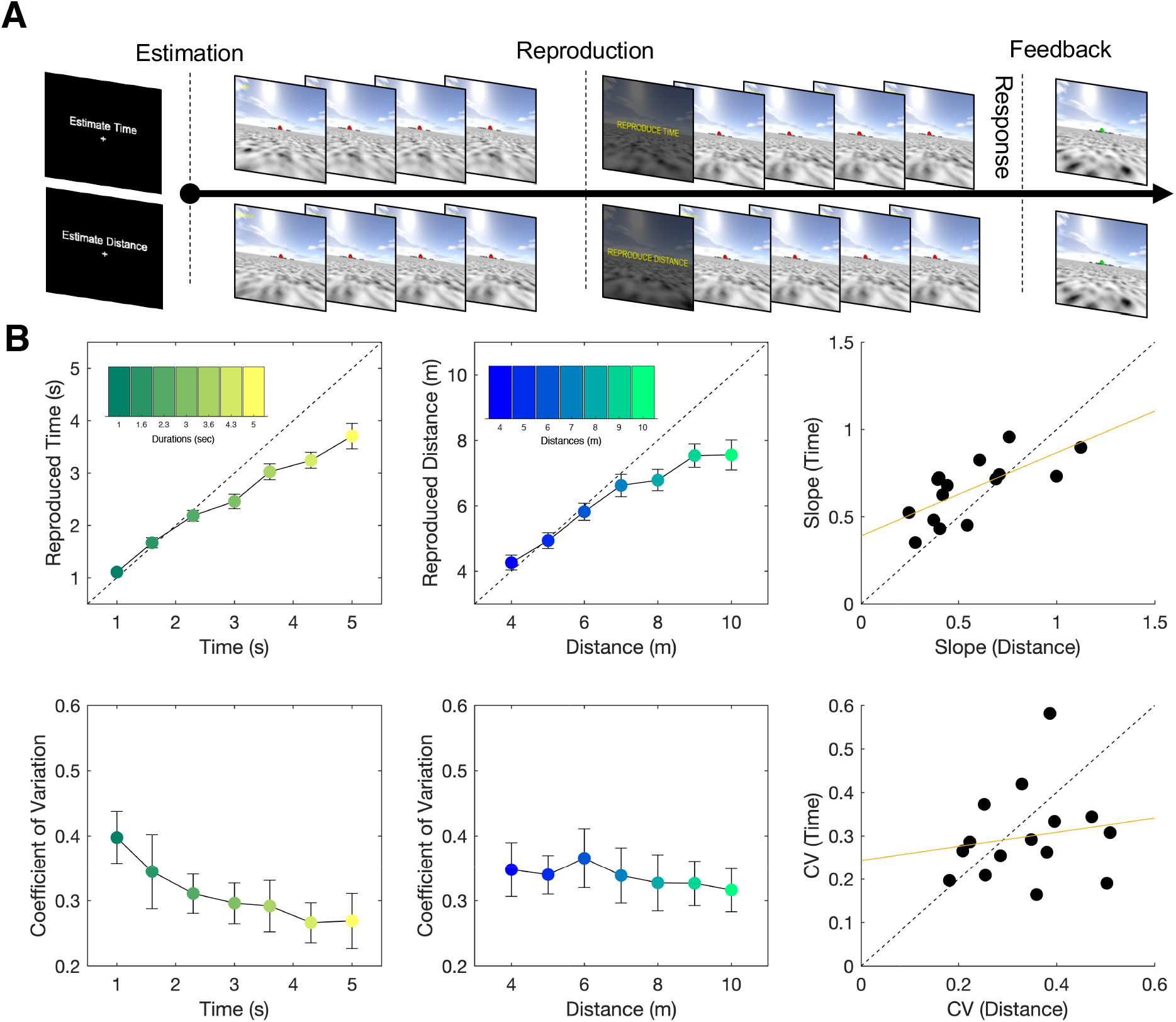
Task and Behavioral Data. (*A*) Subjects performed a distance reproduction task within a VR environment. On a given trial, subjects were initially prompted to estimate either time or distance, after which they were placed at a random point in an open field environment facing a distant red sphere. Subjects moved forwards while estimating the relevant dimension, after which they were told to reproduce that same dimension. Walking speed varied between estimation and reproduction phases. Feedback was provided after subjects made their response. (*B*) Reproduced times and distances as a function of estimated intervals are displayed (Mean *±* SE); subjects exhibited gradual underestimation of both dimensions with longer intervals. Right panel displays slope values for both dimensions, which were significantly correlated; further, subjects exhibited higher slope values for distance reproduction, indicating more accurate performance. Bottom panels display normalized variability (coefficient of variation; CV) for both dimensions for each interval tested. Bottom right panel displays CV values collapsed across interval; no difference or correlation was detected.

### Time and Distance Reproduction activate overlapping networks

In our scanning protocol, we initially performed a univariate analysis of BOLD activation while subjects performed the task. Previously, using the distance version of this task, we observed that the reproduction phase activates a bilateral frontoparietal network of regions, as well as subcortical sites across the basal ganglia and hippocampus (17). Our findings here replicate this effect, with distance reproduction activating a broad network of brain regions across the basal ganglia, cerebellum, anterior cingulate cortex (ACC), right prefrontal cortex (PFC), and parietal and occipital regions, as well as the right hippocampus (Figure 2, Table 2). Notably, time reproduction activated a similar network of regions, although without any activation in the hippocampus (Figure 2, Table 1). In contrasting these effects against one another, we observed a striking dissociation between dimensions across the central sulcus; distance reproduction preferentially activated a variety of posterior regions, including a cluster of regions extending from the middle occipital gyri to the inferior parietal lobes and from the thalamus to the cerebellum, including the bilateral hippocampus (Figure 2, Table 3). In contrast, time reproduction exclusively activated a single region in anterior cortex the bilateral supplementary motor area (SMA; Figure 2, Table 3). These findings suggest a division-of-labor between rostral and caudal parts of the brain in coordinating movements for distance and time.

**Figure 2:**
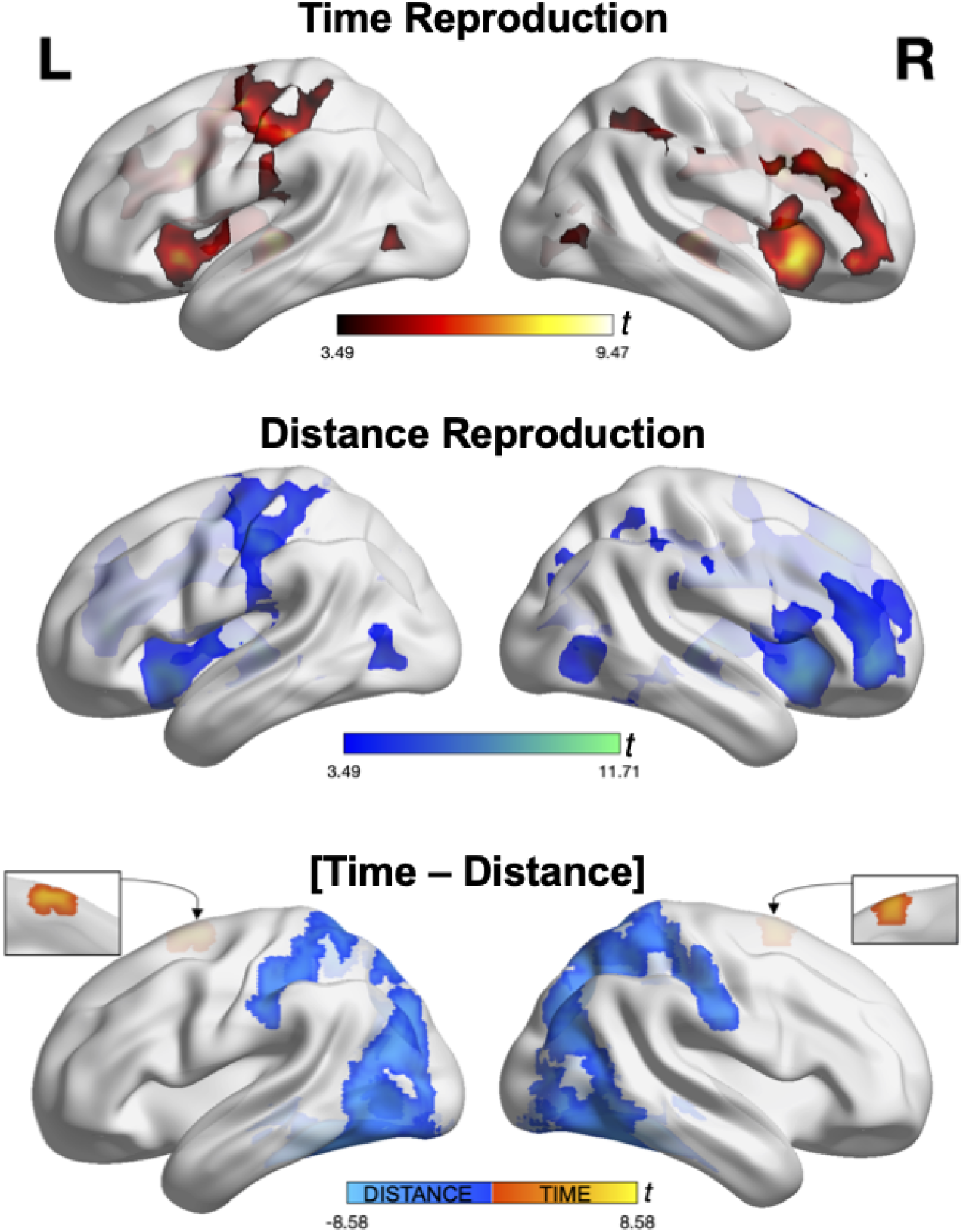
Univariate fMRI results for Time and Distance Reproduction. Top panel displays significant activation when subjects reproduced time estimates. Clusters of activation were observed in frontoparietal structures, including inferior frontal gyrus, inferior parietal lobe, supplementary motor area, basal ganglia, and cerebellum (not shown; see supplementary materials). Middle panel displays significant activation when subjects reproduced distance estimates. Clusters of activation were again found in similar frontoparietal regions, as well as subcortical regions including basal ganglia, cerebellum, and also hippocampus (for full list of activated regions see supplementary materials). Both time and distance contrasts were from comparing reproduction phase activity against estimation phase activity. Bottom panel displays the contrast of time with distance reproduction. Here, distance reproduction preferentially activated posterior regions across striate and extrastriate cortex, as well as bilateral parietal regions and the hippocampus. In contrast, time reproduction only activated the supplementary motor area bilaterally. All clusters thresholded at height p*<*0.001 uncorrected and cluster p*<*0.05 FWE error corrected.

### Time and Distance can be decoded during reproduction

To further examine distance and time estimates in the brain, we employed a multivariate decoding technique, in which a whole-brain searchlight (6mm) was used to train a support vector machine on intervals of time and distance. During the reproduction phase, we again observed a dissociation between decoding accuracy across two distinct regions. For time, the only region where significant decoding accuracy was observed was in the SMA [coordinates]. For distance, in contrast, we observed significant accuracy in several areas, including the left caudate and cerebellum, yet the only region to survive our significance threshold was the retrosplenial cortex (RSC) (Figure 3; Table 4). Examination of confusion matrices for both regions revealed relatively accurate performance (8), although in both maps there was a tendency to underestimate the true interval as they became longer (Figure 3).

**Figure 3:**
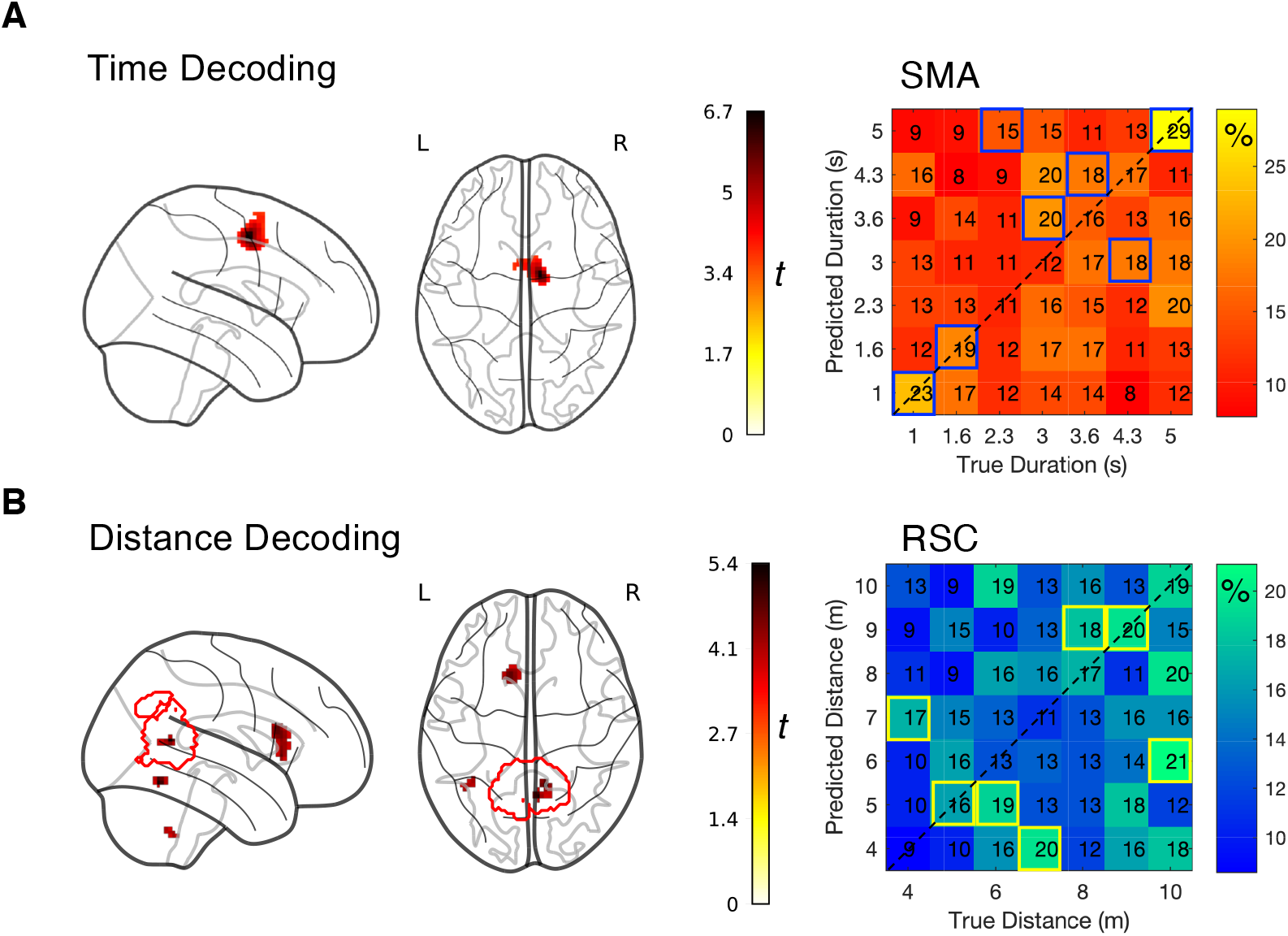
Decoding of Time and Distance Estimates. (*A*) Decoding of time estimates during time reproduction revealed activation only within the supplementary motor area (left panels). Right panel displays the average confusion matrix for this region, containing the percentages of predicted durations against the true duration label; blue squares highlight the maximum predicted duration for each true duration. (*B*) Decoding of distance estimates during distance reproduction revealed significant decoding in several clusters, but with only the retrosplenial cortex passing cluster correction; red contour signifies the mask used for retrosplenial cortex, derived from NeuroSynth. Right panel displays the average confusion matrix for this region, again with percentages of predicted distances against true distances; yellow squares highlight the maximum predicted distance for each true distance. All significant clusters at height p*<*0.001 and cluster p*<*0.05 FWE corrected.

### Multivariate Timing differs between Estimation and Reproduction

To further examine decoding performance, we next examined accuracy in the estimation phase, when subjects walked for an unknown distance or time. Here, no significant regions were detected for the distance estimation phase. However, the time estimation phase exhibited significant accuracy decoding in several regions, including the SMA, precentral gyrus, and right inferior frontal gyrus (rIFG). When examining the SMA cluster further, we again observed in the confusion matrices a tendency for the classifier to underestimate the true duration (Figure 4). As a comparison with the reproduction phase, we extracted the beta values from the SMA region for estimation and reproduction phases and compared them. Recent work has suggested that neurons in the SMA complex exhibit tuning properties for duration, such that distinct sub-regions preferentially activate for longer intervals in a gradient fashion (34; 35). On the other hand, the SMA region has also exhibited activation patterns consistent with a temporal accumulator, in which activity increases with elapsing intervals (36). When examining the beta values for each interval, across estimation and reproduction phases, we observed that beta values were flat while estimating progressively longer durations, consistent with a tuning mechanism account (8). However, during the reproduction phase, we instead observed a linear increase in the modeled BOLD response in the SMA (Figure 4). A repeated measures ANOVA confirmed these observations, with a significant effect of phase (estimation vs reproduction [*F* (1,15)=15.896, *p*=0.001, *η*^2^=0.217]) and phase by interval interaction [*F* (6,90)=3.06, *p*=0.009, η2=0.043]. As such, both patterns were observed in the SMA at distinct phases, suggesting the SMA operates in two modes when timing virtual movement, a point we turn to in the discussion.

**Figure 4:**
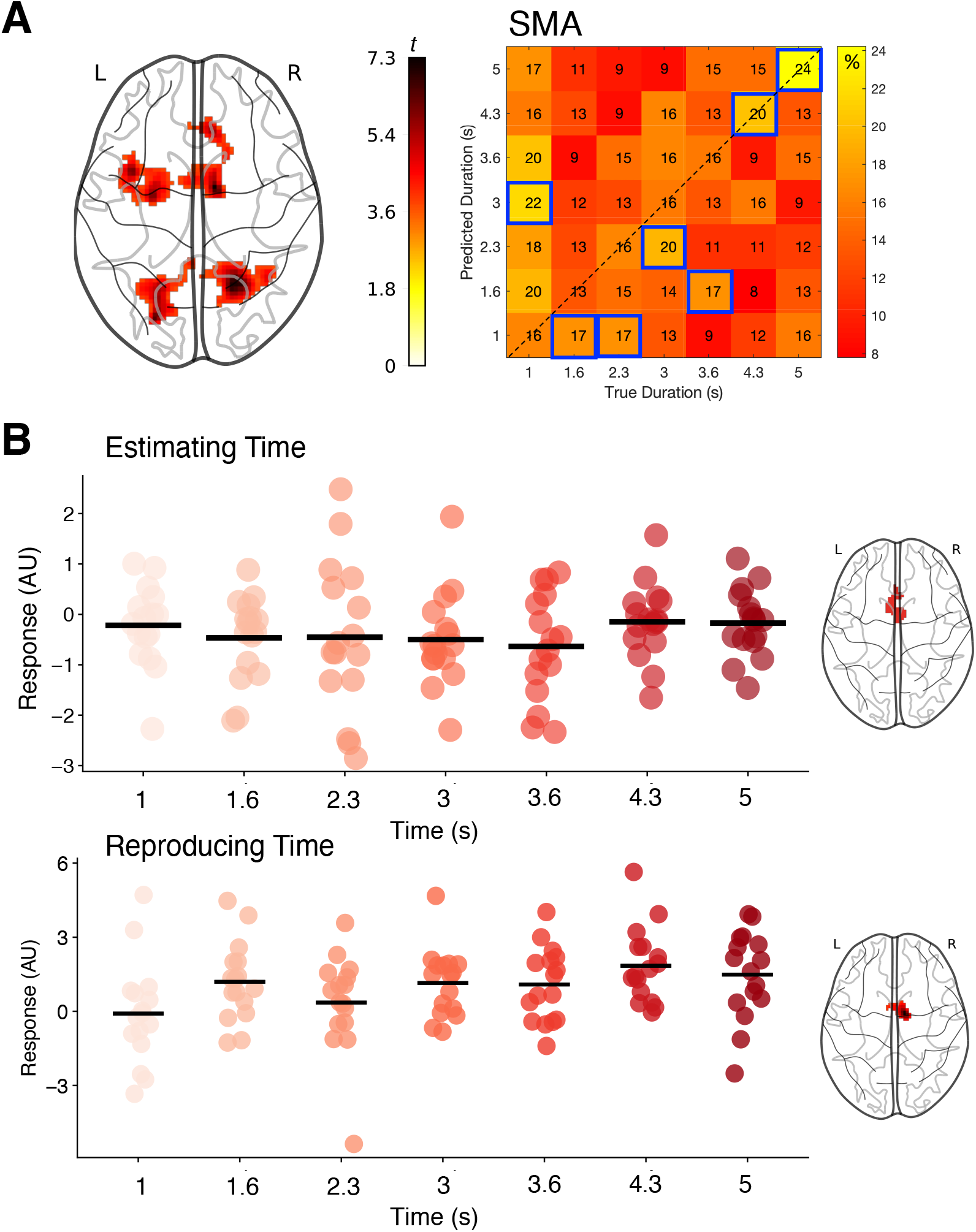
Comparison of time decoding in the supplementary motor area reveals different coding properties for estimation and reproduction. (*A*) Significant clusters for decoding time walked during the estimation phase, revealing activity in supplementary motor area, precentral gyrus, and bilateral superior parietal cortex. At right, the average confusion matrix for the SMA region, displaying percentage of predicted duration against true duration; blue squares indicate maximum predicted durations for each true duration. (*B*) Beta values within the SMA region (averaged across regions from right brain images) for the time estimation and reproduction phases; black bars indicate the mean. For the estimation phase, beta values exhibited no difference across presented times. However, for the reproduction phase, SMA beta values increased linearly with increasing duration. All clusters significant at height p*<*0.001, clusters p*<*0.05 FWE corrected.

### Cross-classification decoding

As a final comparison, we tested for abstract representations of magnitude. To accomplish this, we took a cross-classification approach (37), in which a classifier was trained on activation patterns from the seven intervals in one dimension (e.g. time) and tested on patterns from intervals in the other dimension (e.g. distance). Previous work has demonstrated that the right inferior parietal lobe may serve as an abstract representation of magnitude, such that symmetrically longer and larger stimuli are represented in this region, regardless of dimension (4). We examined classification accuracy bidirectionally, such that significant performance represented regions that could cross-classify above chance in either direction. Here, two regions were observed that survived significance testing: the right supramarginal gyrus (rSMG) and a cluster within the right middle occipital gyrus (Figure 5; Table 4). To further determine the nature of the occipital cluster, we compared this region to a recently released probabilistic functional atlas of visual cortex (38). Here, we observed that the occipital cluster most closely overlapped with the probable location of the occipital place area (OPA), a region that activates in response to visual scenes, with potential navigation functions (39).

**Figure 5:**
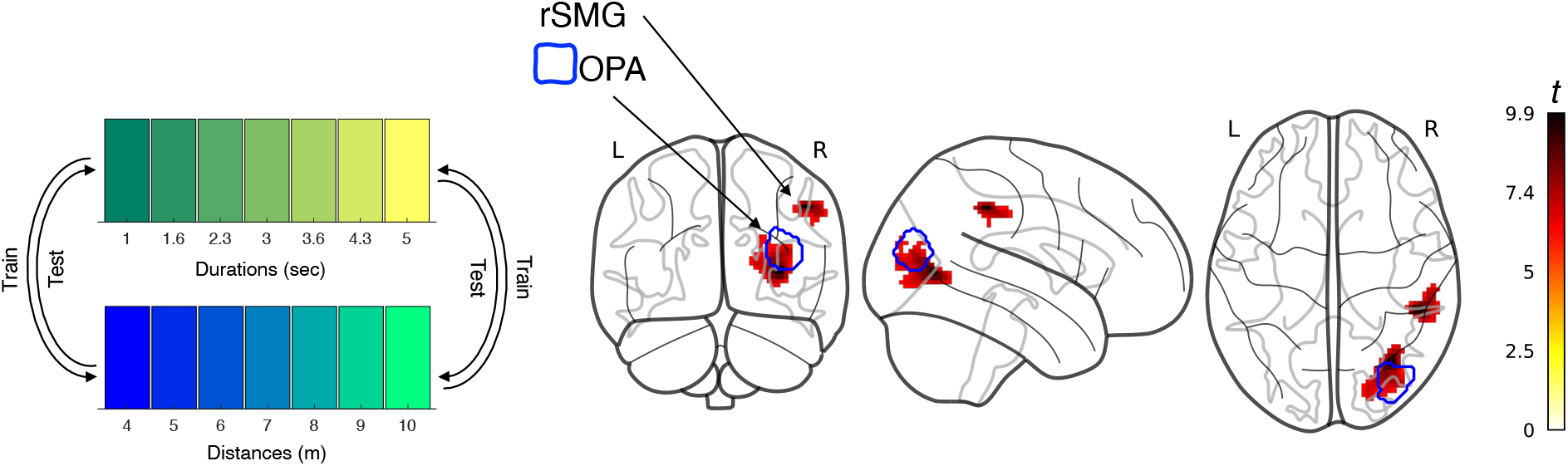
Results of cross-classification decoding, demonstrating amodal representations of time and distance. At left, classifiers were separately trained one dimension (e.g. time) and then tested on the alternate dimension (e.g. distance). At right, two significant clusters of activation were observed within the right supramarginal gyrus and lateral occipital cortex. Blue contour represents a probable mask of the occipital place area, derived from (38).

## Discussion

Behaviorally we observed central tendency for both time and space, similar to other studies using the same paradigm; however, this effect was significantly larger for time than for space and was correlated between dimensions. This suggests that there may be a common mechanism for reproducing different magnitudes regardless of dimension (i.e., time, space, size, etc.). On the other hand, variability of reproductions showed no difference between time and distance; therefore, participants were equally consistent when reproducing either dimension.

Similar to previous fMRI findings (17), during distance reproduction a broad network of brain regions were activated including the basal ganglia, cerebellum, ACC, rPFC, parietal and occipital regions, and the hippocampus. Additionally, during time reproduction similar networks were activated excluding the hippocampus. When contrasting the activation during reproduction between both dimensions we found a dissociation across the central sulcus in that, distance activated a cluster of posterior regions extending from the middle occipital gyri to the inferior parietal lobes and from the thalamus to the cerebellum and bilateral hippocampus; whereas time solely activated the SMA, bilaterally. This suggests that the brain coordinates movements for distance and time between rostral and caudal regions. We additionally found that time and distance can be decoded during reproduction; specifically, the region that could most accurately decode time was observed in the SMA whereas the region that could most accurately decode space was observed in the RSC.

We additionally examined decoding performance for the estimation phase of the task, in this case no specific regions could accurately decode the distance estimation phase; however, several regions could decode the time estimation phase including the SMA, precentral gyrus, and rIFG. Given that the SMA was able to decode time in both estimation and reproduction we further examined this region by extracting beta values for both phases and compared them. We found that when participants were estimating longer durations the beta values remained flat suggesting a tuning mechanism account(34; 8); however, in the reproduction phase we observed a linear increase in the BOLD response in the SMA. This finding suggests that the SMA operates in different modes when timing virtual movement depending on the phase (i.e., estimation vs reproduction). The estimation phase finding suggesting a tuning mechanism account is in line with previous work which suggests that the SMA is chronotopically organized in a rostrocaudal gradient in which mor(23)e rostral neurons encode shorter durations and more caudal clusters encode longer durations (34; 35). Whereas the reproduction finding is in line with work which suggests that the SMA exhibits activation patterns consistent with a temporal accumulator, similar to properties of the CNV during EEG, in which activity increases with elapsing intervals (36). We further note that the linear activation pattern of the SMA is timelocked to the *onset* of the reproduction phase. This pattern shows a striking similarity to that observed recently in EEG data from this same task (21); larger CNV amplitude responses were observed corresponding to longer *intended*, or planned intervals. Considering putative correspondence between the CNV and SMA activity (23), we suggest both reflect the same mechanism. More specifically, we suggest this amplitude change reflects a “pre-planning” signal for reproducing time intervals, as has been observed previously in non-human primates (40).

Lastly, we wanted to further corroborate previous work suggesting that the right inferior parietal lobe is important for the representation of different magnitudes regardless of dimension using a cross-classification approach (4). We note that two regions were found to cross-classify above chance in either direction, the rSMG located in the right inferior parietal lobe as well as the right middle occipital gyrus. The rSMG finding suggests that the right inferior parietal lobe does represent general magnitudes. We found that the occipital cluster appeared to overlap with the probable location of the OPA which activates when viewing visual scenes and has been suggested to be responsible for potential navigation functions (39; 38).

Taken together these findings suggest that reproducing time and distance activate a similar corticalsubcortical network of regions divided between anterior and posterior regions, respectively, and that the duration and distance walking can be accurately decoded from activation in the SMA and RSC, respectively. Additionally, the findings support a potential amodal decoder in the rSMG and occipital cortex located approximately at the OPA suggesting this area of the brain encodes both distance and duration traveled from a starting location in the environment. Further, the findings point to a simple circuit model describing how time and distance measurements interact (Figure 6). Specifically, duration and distance estimates appear to be encoded via separate channels, each of which may be manipulated by the amount of attention focused on them; these estimates then interact in memory, prior to judgments being made at a planning stage. Neurally, the RSC, SMA, IPL, and prefrontal cortex (PFC) all appear to be active, but in different ways. Specifically, the RSC and SMA both appear to encode and store intervals of distance and duration, respectively. These intervals are then passed to the PFC as vectors for movement (Harootonian, et al. 2021; Leek, et al. 2008) that include both the distance (*V* _*d*_) and duration (*V* _*t*_) of the required action. Further, we suggest the IPL encodes the covariance of distance and time as a supramodal signal for magnitude.

**Figure 6:**
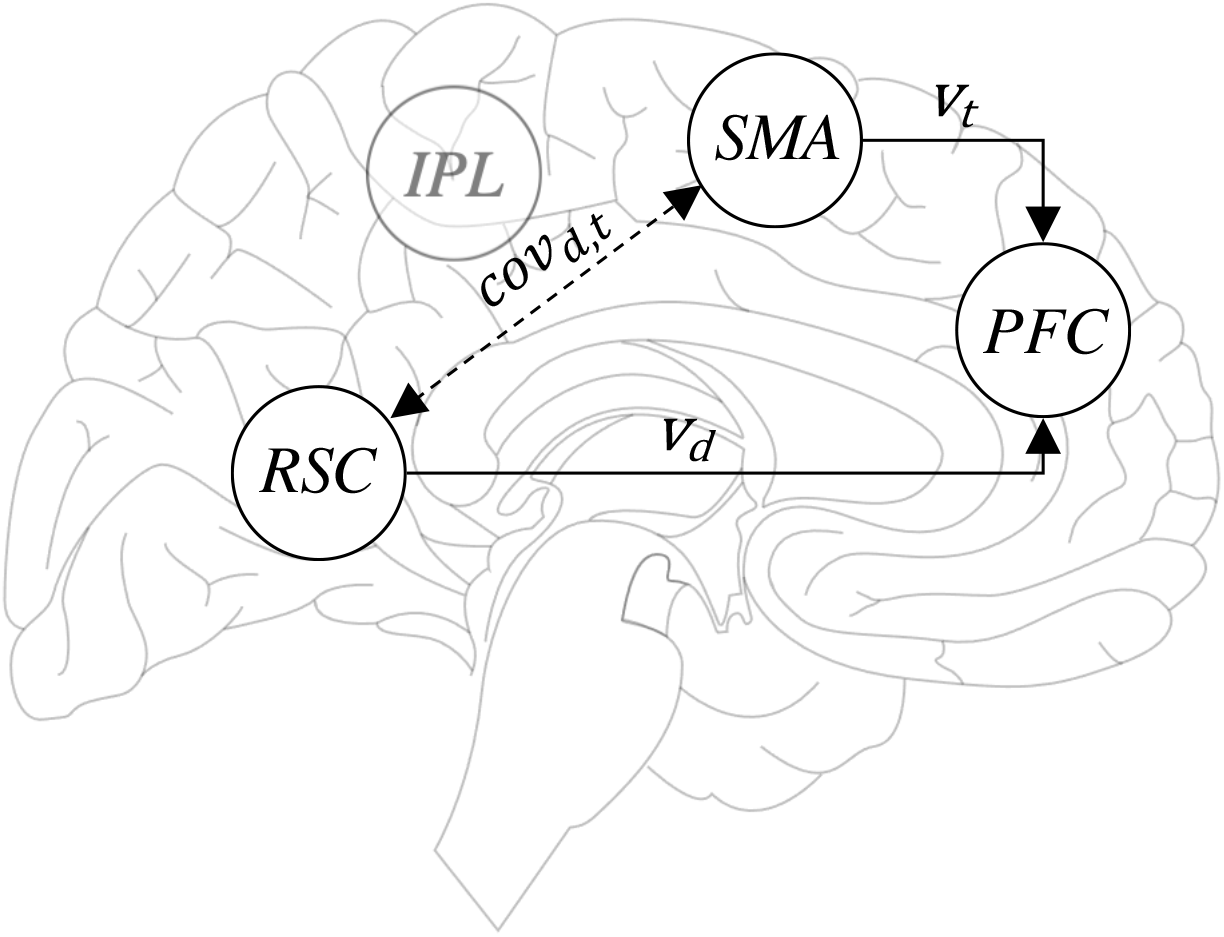
Diagram of a simple model of results. The SMA and RSC serve as reference points for time and distance, respectively. Accordingly, each region encodes a vector (*v*), for time (*v*_*t*_) in the SMA and distance (*v*_*d*_) in the RSC while navigating from an origin point. These vectors are in turn processed by prefrontal regions during path integration. Additionally, the covariance between distance and time (*cov*_*d,t*_) is implicitly encoded in the IPL, which shares connections with each of these regions.

**Figure 7:**
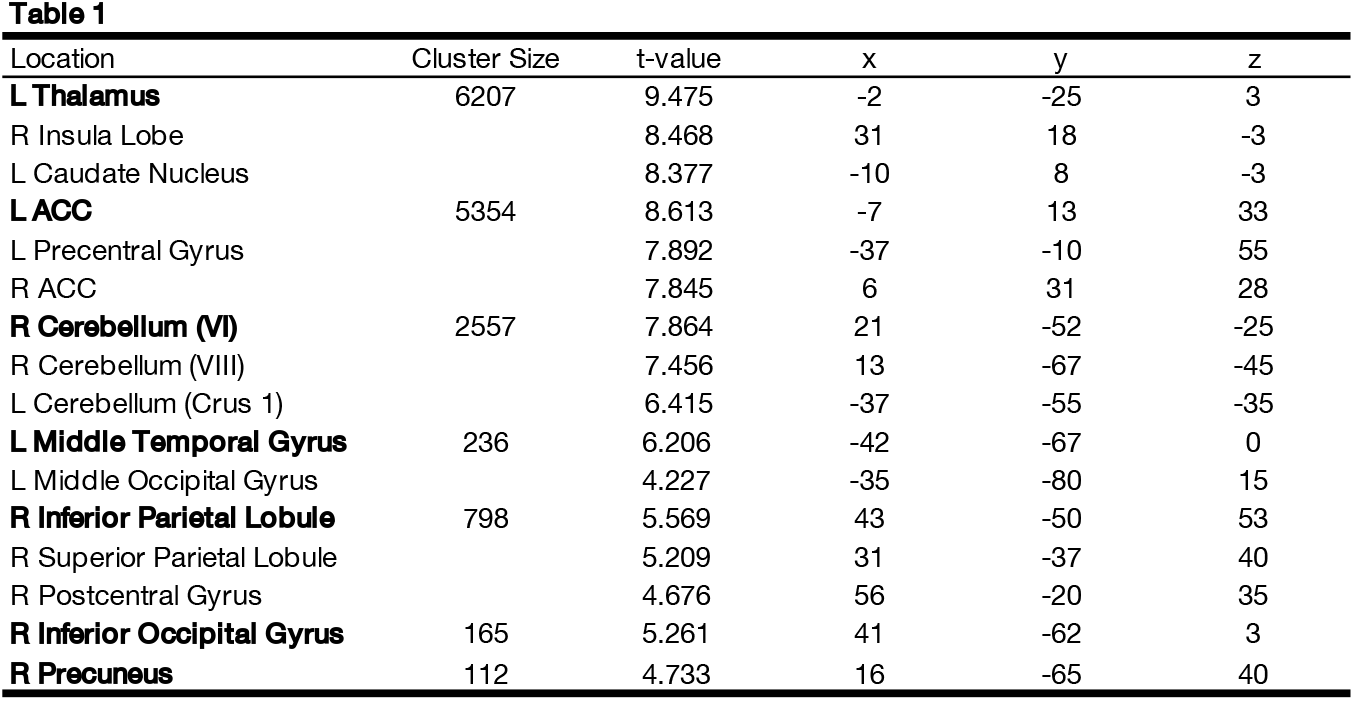
Table 1

**Figure 8:**
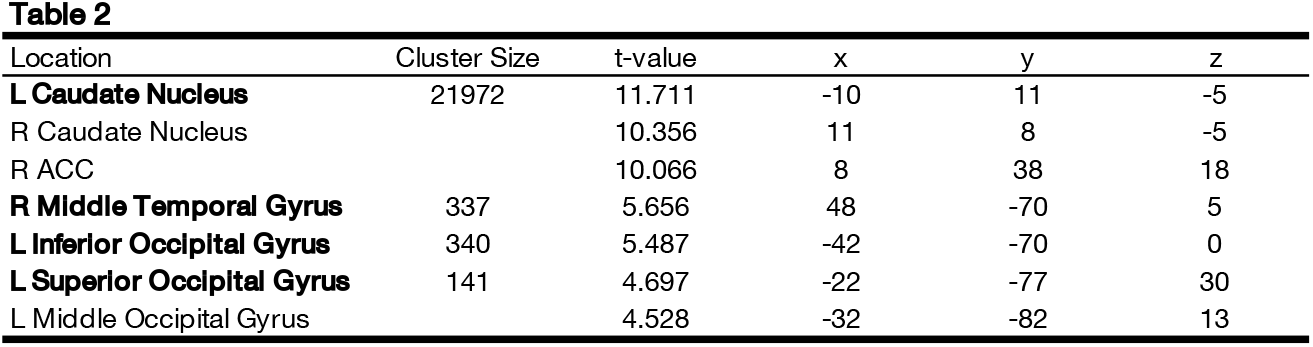
Table 2

**Figure 9:**
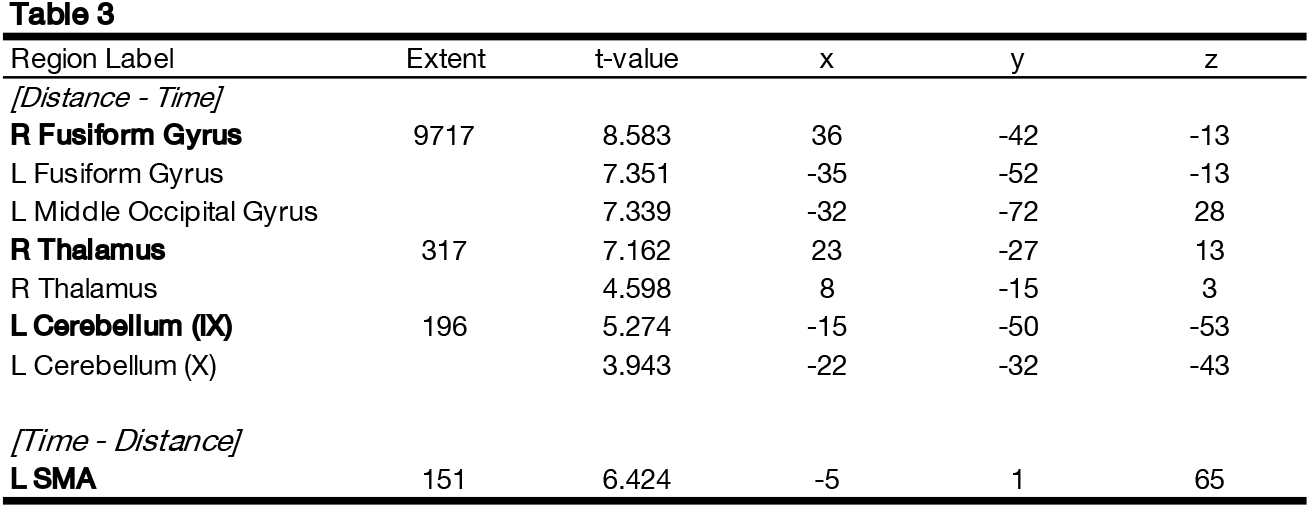
Table 3

**Figure 10:**
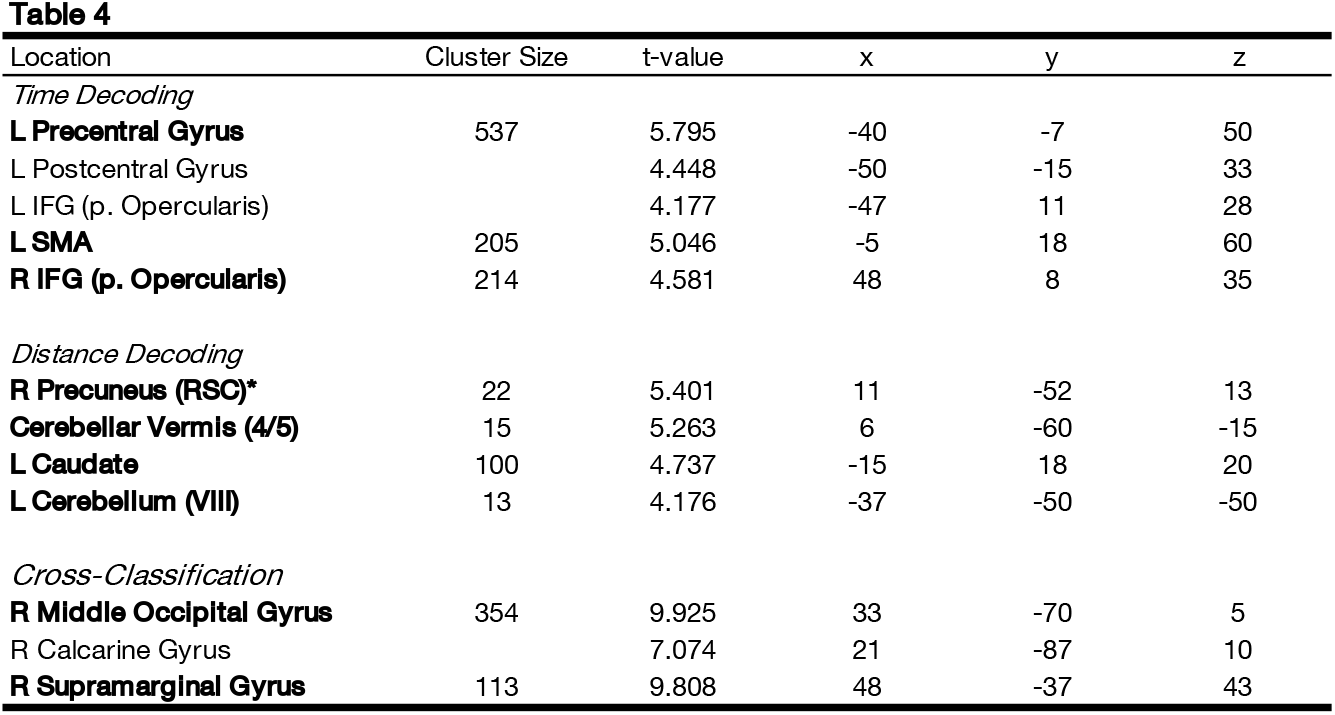
Table 4

We suggest this circuit model provides a parsimonious explanation for the roles of the RSC and the SMA in the processing of time and space. For space, the RSC has been associated with path integration, transitions between viewpoints, and landmark recognition (41). In particular, the RSC may serve as an anchor for movements in the environment, such that an observer can keep track of their point of origin (42). For time, the SMA has been associated with a temporal accumulation process, but also has shown involvement in tracking relative differences between experienced time and remembered intervals (43). We suggest that this latter comparison process is similar to placing an anchor in time; that is, tracking how much time has elapsed from a given event, a necessary component of navigating through the environment. Both processes may be deployed in the service of action, for choosing between different paths to travel to achieve a particular goal.

Overall, our findings demonstrate that time and distance can be both separately, and conjointly, decoded from brain activation patterns in distinct parts of the brain. These findings suggest that the brain holds separate metrics for each dimension, while also tracking a general magnitude for travel. Future studies will be necessary to determine how time and space metrics are combined downstream, and how disruptions in one dimension may impact the processing of the other.

## Materials and Methods

### Participants

This study included 23 right-handed participants that were neurologically and psychiatrically healthy (15 females, 7 males, 1 undisclosed; 18-29 years old, M = 22.2) recruited at George Mason University. All participants satisfied the MRI safety screening criteria and provided consent as approved by the University Institutional Review Board (IRB) at George Mason University. Participants were compensated for their time with monetary payment. Due to a software error during data collection, we note that 7 subjects did not have their behavioral data saved; as such, all behavioral and multivariate decoding analyses are conducted on the remaining 16 subjects, whereas the univariate analyses are reported on the full 23 subjects.

### Task

Participants completed temporal and spatial reproduction tasks in a virtual environment (VE) while undergoing functional magnetic resonance imaging (fMRI) using a SIEMENS Prisma 3T scanner with a 32-channel head coil. The VE was previously used in behavioral (3) and EEG (21) studies from our lab. The paradigm was originally developed by (15) and designed using Vizard 5.0 (Worldviz), a Python-based software. The ground in the VE was created using a black and white noise image that resembled a textured “desert” ground. There were 20 identical scattered rocks imported from SketchUp 3D (Trimble Navigation) and a sun, in the form of a sphere on the horizon. The clear sky was a simulated 3D dome included in the Vizard software. Importantly, the construction of the VE minimized reliable environmental distance cues by randomizing the initial viewpoint and position/orientation of each rock at the start of each trial. Additionally, the 3D sky utilized made it appear as though the horizon was always a constant distance away. Participants completed the task by viewing the VE through a head coil mounted mirror which reflected the monitor (Cambridge Research Systems Display++, 120Hz refresh rate) at the end of the scanner’s bore during scanning and respond using an MR compatible handheld button box (Current Designs) with four buttons aligned horizontally (only furthest left and right buttons used). Procedure: Trials were completed in blocks of 14 trials per block, for eight blocks, each lasting approximately 6 minutes. TIME trials and DISTANCE trials and were presented randomly in within each block, consisting of 7 possible trials of each type. Each trial consisted of an estimation and reproduction phase. The estimation phase began after a white fixation cross on a black background for a variable interval drawn from an exponential distribution with a minimum duration of 3 s (17); above the fixation cross subjects viewed the words “Estimate Time” or “Estimate Distance”. Subjects began with a random view of the horizon in the VE and a red sphere appearing on the horizon. Participants pressed the left button to begin movement toward the red sphere. After a particular distance (DISTANCE trials) or temporal interval (TIME trials) had passed, movement was automatically stopped, the screen dimmed and the words “REPRODUCE DISTANCE/TIME” appeared. Following a variable 4-8s delay (uniform distribution) the words disappeared, the screen re-illuminated, and the reproduction phase began. The reproduction phase began the same as the estimation phase (left button press to begin movement) and participants reproduced the distance or time traveled in the estimation phase by terminating the movement with a right button press. As a reminder, the appropriate magnitude was displayed in the upper left corner (e.g. the word “TIME” was displayed on time trials) to ensure that subjects did not forget the dimension to be reproduced (18). The sphere on the horizon provided feedback by appearing green if reproduction was accurate or red if it was inaccurate. The feedback was adaptive on a trial-by-trial basis such that the feedback constant (k) was updated in a 1-up/1-down rule (step size = 0.015) if the reproduced interval was within (step-down) or outside of (step-up) the feedback window (interval/k). Starting k value was 3.5 (44; 21) and was calculated separately for time and distance. The distances used in the estimation phase of DISTANCE trials was randomly selected from seven linearly-spaced intervals between 4-10m. On the estimation phase of TIME trials the distance was determined by the temporal interval and varied across seven linearly-spaced intervals between 1-5s. The speed of movement was randomly selected from a uniform distribution between 2 and 4.5 m/s in order to match the duration experienced on spatial reproduction trials to those presented in temporal reproduction trials (21). Additionally, in order to eliminate the possibility of participants using the time spent moving as a measure of distance or the distance walked as a measure of time traveled, the simulated walking speed was randomly altered between estimation and reproduction phases such that it was noticeably faster or slower than the estimation phase speed (maximum +60% estimation speed, drawn from a normal distribution). Participants were instructed not to count or tap during the task (45) and were not aware of the range of distances or times but were aware that the walking speed was altered.

### fMRI

Scanning was conducted using a 3T Siemens Prisma Magnetom scanner. All subjects initially received a high resolution, T1-weighted 3-D magnetization prepared rapid gradient echo scan (192 coronal slices, TR = 2400 ms, TE = 2.28 ms, TI = 1060 ms, matrix size 300×320, 0.8 mm isotropic voxels). Following this, a gradient echo fieldmap was collected (64 coronal slices, TR=1200ms, TE=33ms, matrix size 84×84, voxel size 2.5mm isotropic voxels). Multiband gradient-echoplanar images (EPI) were individually acquired (64 coronal slices, TR = 1200 ms, TE = 33 ms, matrix size 84×84, voxel size 2.5 mm isotropic voxels). EPI volumes were acquired in eight separate runs, with each run lasting a variable duration (6 min). The first three volumes of each run were additionally discarded to allow for steady-state magnetization.

### Behavioral Analysis

Behavioral data were analyzed similar to our previous report (21). Briefly, the mean reproduced duration and distance were calculated for each subject and interval tested. Additionally, the coefficient of variation (standard deviation / mean) was calculated as a measure of variability. To further quantify performance, we fit mean reproductions with a simple linear regression and calculated the slope values for each dimension; a slope value closer to zero indicates a larger central tendency effect, wherein responses gravitate to the mean, while a value closer to one indicates more veridical performance.

### fMRI Analysis

Analysis of fMRI images was conducted using SPM12 (https://www.fil.ion.ucl.ac.uk/spm/). Standard preprocessing steps were conducted for each subject, including realignment, normalization, and smoothing (6mm). Additionally, prior to realignment we corrected for fieldmap homogeneity using Hyperelastic Susceptibility Artifact Correction (HySCO) (46) module in the ACID toolbox for SPM. At the first-level (single-subject), a general linear model was constructed by convolving the onset times of events with a canonical hemodynamic response function (hrf). More specifically, we specified events at the onset of either the estimation phase or the reproduction phase of time and distance trials, separately. Additionally, we modeled the hrf with a boxcar function, in order to account for the varying durations of each interval (17). At the second-level (group) we conducted several planned contrasts, including [Time Reproduction *>* Time Estimation], [Distance Reproduction *>* Distance Estimation], and [Time Reproduction - Distance Reproduction]. Note that the final contrast was examined in both directions. For this and all other comparisons, significance was set at a voxel level threshold of p*<*0.001, and a cluster-level threshold of p*<*0.05, familywise error (FWE) corrected. Anatomical localization was conducted using the SPM Anatomy Toolbox. To further interrogate activity within the RSC, we defined a functional map using the Neurosynth database (www.neurosynth.org) with the search term “retrosplenial cortex”; a forward inference z-map was extracted and thresholded at z=2.5 to include only voxels within the precuneus region, behind the posterior commissure (41). This map was converted into a binary mask that was used in all analyses with a small volume correction.

Multivariate decoding was performed using The Decoding Toolbox (47). To perform this, we used functional data from the step prior to smoothing, in order to maintain more granular activation patterns. A new firstlevel GLM was constructed, with separate regressors for each interval of time and distance tested. These resulting beta maps were subjected to a whole-brain searchlight analysis (6mm searchlight) in which, at each location a support vector machine was trained and tested in a leave-one-run-out cross-validation procedure on the seven intervals tested for each dimension. Separate decoding analyses were conducted for time and distance, at both the estimation and reproduction phases, resulting in [accuracy-chance] maps for each condition. These maps were then smoothed as for the univariate analysis with a 6mm Gaussian, and then combined for a group-level analysis using one-sample t-tests with the same significance threshold as above. Cross-classification analysis was conducting using the `make_design_xclass` function from The Decoding Toolbox, in which the intervals for time and distance were randomly swapped, again in a leave-one-run-out cross-validation procedure.

